# Epigenetic mediated functional reprogramming of immune cells leads to HBsAg seroconversion in Hepatitis B Virus Reactivation patients

**DOI:** 10.1101/2023.08.21.554133

**Authors:** Jayesh Kumar Sevak, Mojahidul Islam, Gayantika Verma, Anoushka Saxena, E Preedia Babu, Shahana Parveen, Ankur Jindal, Manoj Kumar Sharma, Gayatri Ramakrishna, Shiv Kumar Sarin, Nirupama Trehanpati

## Abstract

**Background:** Hepatitis B virus (HBV) modulates epigenetic landscape by epigenetic regulators. HBsAg seroconversion is possible with immune activation, therefore we aimed to investigate epigenetic modulation in HBV reactivation (rHBV) for viral clearance and seroconversion.

**Methods:** Sixteen retrospectively collected rHBV patients [Seroconverters (SC, n=7, HBsAg loss and anti-HBs>10 IU/ml), non-seroconverters (NSC, n=9)], chronic hepatitis B treatment naïve (nCHBV, n=7) patients and healthy controls (HC, n=7) were included in this study. Genome methylation, gene expression, plasma-cytokines, and immune cell profiling was analysed by Reduced Representation Bisulfite Sequencing (RRBS), QRT-PCR, multiplex-cytokine-bead array and flow-cytometry.

**Results:** rHBV patients having high HBV DNA and ALT showed epigenetic remodellers; KDM2B, NCOR2 and GATA6, immune and metabolic genes; TGF-β, IL-6, IRF8, RPTOR, HK3 significantly (p<0.05) hypomethylated at specific CpG islands compared to nCHBV. TOX was hypomethylated in nCHBV suggesting immune-exhaustion. At-baseline, seroconverters showed hypomethylation of KDM2B, COX19, IRF8, TLR5 and hypermethylation of LAG3 compared to non-seroconverters. Further, in seroconverters at week-24, IL17RA, IFN-γ, TGF-β, and STAT5B (p<0.05) were additionally hypomethylated at specific CpG islands suggesting immune activation. Cytokine-bead analysis revealed increased IL-6 (p=0.009) and decreased LAG3 plasma levels (p=0.01) also imply on significantly differentiated HBV specific CD8, Tfh and Th1/17 cells in seroconverters at baseline and week-24. However, both nCHBV and non-seroconverters had consistent hypomethylation of LAG3 and TOX, which leads to immune exhaustion.

**Conclusion:** In rHBV, seroconversion is driven by position specific CpG islands methylation in epigenetic remodellers, immune and metabolic genes. Immune metabolic reprograming is reflected by Th1/17 differentiation, extensive interleukin production for HBsAg seroconversion.

**Graphical Abstract:** 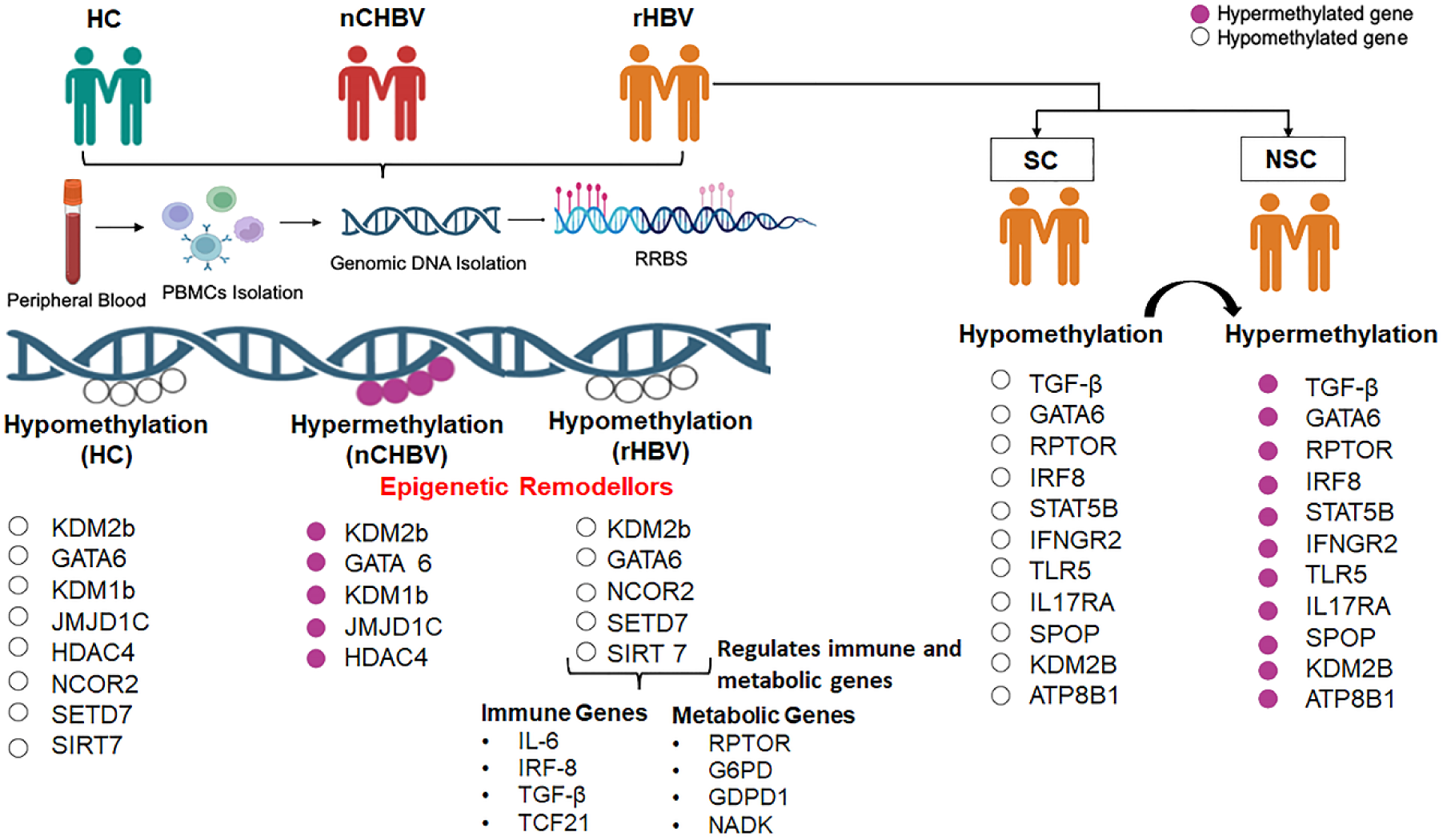

**Lay summary:** Epigenetic landscape in nCHBV depicts exhaustion and immune dysfunction. Out of many hypermethylated CpG islands of nCHBV, few become hypomethylated in rHBV and drives immune and metabolic reprogramming. This study provides insights into the cellular and molecular basis of epigenomic programs that regulate the differentiation and activation of immune cells leading to viral clearance and seroconversion. Targeting epigenetic mechanism could be promising strategy for the treatment of nCHBV and non-seroconverters.

## Introduction

The hepatitis B virus (HBV) infection represents a global public health concern with significant morbidity and mortality. Globally, 296 million people are estimated to be chronically infected with HBV.[1] Despite the existence of effective prophylactic vaccines and approved antiviral regimens, such as immune modulator PEG-IFN (polyethylene glycol interferon) and nucleos(t)ide analogues (NAs), which efficiently suppress viral replication, resolution of infection is rarely achieved. [2–3]

Functional cure is defined as loss of HBsAg and detection of anti-HBs (>10mIU/ml) in a person who was previously HBsAg positive and anti-HBs negative.[4] Coordinated crosstalk between adaptive and innate immune system can accomplish an effective viral clearance. However, there is a probability of HBsAg clearance (1 to 2%) with or without the appearance of anti-HBs titres (>10 IU/ml) with immune activation in reactivation.[5] HBV clearance is mainly dependent upon HBV specific CD8^+^ and CD4^+^ T cells. CD4^+^ T cells render help to develop long-lasting effector CD8^+^ T cells, which produce an array of cytokines including IFN-γ and TNF-α responsible for HBV inactivation in the target cells. Blockade of IFN-γ and TNF-α also abrogated the noncytolytic inhibition of HBV. However, impaired T cell responses were observed in chronic hepatitis B patients. [6,7]

Epigenetic profiling of primary CD8^+^ T cells demonstrated the role of histone modifications, DNA methylation, chromatin accessibility in T cell differentiation, activation, memory and effector response during viral infections. [8,9]

Plasticity of T helper (Th) subsets (Th1, Th2 and Th17) and Tregs cells contribute to effector immune response which could also be modulated by epigenetic modifiers.[10] Histone deacetylation and DNA demethylation increase the stable expression of forkhead box P3 (FOXP3) and strengthen the suppressive function of Treg cells. [10] Epigenetic modulation reduces the FOXP3 binding to interleukin (IL)-2, tumor necrosis factor (TNF)-α, and interferon (IFN)-γ promoters in lentivirus-specific CD8^+^ T cells. [11] At the same time, with epigenetic modulation, reduced expression of co-stimulatory molecules like CD40, CD80, CD86 and secretion of pro-inflammatory cytokines including TNF-α, IL-1β, IL-6 and IL-12 inhibits the activation of Th1 and Th17 responses and granzyme B producing CD8^+^ T cells. [12,13]

However, recently epigenomic and transcriptomic regulome revealed that HBV-specific CD8+ T cells which are not terminally exhausted showed long-term memory and polyfunctionality in chronically HBV-infected HCC patients. [14]

Indeed, 54% of the HBV-specific T cells had effector and memory status with increased accessibility of JUN, FOS, BACH1/2, and NFE transcription factors in the CCL5, CD28 and CXCR4 loci. In addition, 13% and 9% of HBV-specific Tregs and Tex cells, respectively were regulated by a unique and shared program of the NFKB1/2-, REL-, NFATC2- and NR4A1- TCR associated epigenomic regulome. [14]

Furthermore, expression defects of histone lysine demethylase (KDM) also have important role in immune dysfunction. KDM demethylases induce immunity by modulating H3K9 methylations. [15] Inhibition of KDM4, impairs the glycolysis in β- glucan-induced trained immunity suggesting a link between alterations in epigenetic changes and cellular metabolism.[16] Innate immunity is also regulated by lysine-specific demethylase 2b (KDM2B), which selectively demethylates H3K4me3 and H3K36me2 in macrophages and dendritic cells for IL-6 secretion but not TNF-α, IL-1, and IFN-β. [17–19]

Therefore, understanding the epigenetic remodellers mediated immune reorganization in HBV reactivation (rHBV) for viral clearance and seroconversion is very important for future immune therapies. In this study, we focused on site specific methylation of epigenetic remodellers, immune and metabolic genes involved in regulation of the T-cell differentiation and cellular metabolism for CHBV persistence, viral clearance and seroconversion in HBV reactivation patients.

## Materials and Methods

### Clinical characteristics of patient’s enrolment

Retrospectively collected sixteen HBV reactivation (rHBV) patients [seven seroconverters (SC) and nine non-seroconverters (NSC)], seven naive chronic hepatitis B infected (nCHBV) patients and 7 healthy controls were subjected to methylation studies (Study cohort Fig. S1). SC are characterised with HBsAg loss and development of anti-HBs titres (>10 IU/mL) within 48 weeks.

### Reduced Representation Bisulfite library construction and sequencing

8-10 ml peripheral blood was processed for PBMCs isolation. From PBMCs, we have isolated DNA for preparing Reduced Representation Bisulfite Sequencing (RRBS) libraries using Zymo-Seq RRBS Library Kit (Zymo Research, USA, #D5461). In brief, 300ng of genomic DNA was purified with a GeneJET Genomic DNA purification kit (Thermofisher, USA, #K0721) and digested with restriction enzyme MspI (5’C’CGG) by incubating at 37°C for 4 hrs (Details in Supplementary data). [20]

### RRBS Sequencing Data processing and Analysis

RRBS analysis of rHBV, nCHBV and HC FastQC (version 0.11.8, www.bioinformatics.babraham.ac.uk/projects/fastqc/) was used for quality check. Universal adapter was removed using trim galore (version 0.6.2, https://www.bioinformatics.babraham.ac.uk/projects/trim_galore/), a wrapper script to automate quality and adaptor trimming as well as quality control. Genome indexing was done using Batmeth2. These reads were aligned to the reference Hg19 genome downloaded from Ensemble release 102. Mapping and alignment were performed using Batmeth2 using align tool. Also, DNA methylation level was calculated using Batmeth2 calmeth tool. Then annotation was done using MethyGff tool using hg19 genome gff and genome methylation obtained using calmeth tool. Methdiff.py from BSMAPZ tool (fork of BSMAP V2.90) was used for differential methylation ratio (DMRs) analysis. Selected gene methylations were fetched using perl and shell script.[21] We have identified the immune genes using an integrated analysis platform, innateDB database. Similarly for epigenetic genes we used publicly available epigene database. Further, we have assessed Transcription factor binding sites by using TRANSFAC 4.0 software and the AliBaba 2.1 database.

### Methylation calculation and Analysis

Methylation ratio was calculated by Batmeth2 calmeth tool.[22–24] (Details in Supplementary data).

## RESULTS

### Demographics and clinical parameters

We have randomly recruited age and gender matched 16 rHBV, 7 nCHBV and 7 HC from previously recruited cohort (Islam et al., AP&T 2022) for methylation and gene expression studies (Fig. S1). rHBV patients at baseline were characterized by ALT>5xULN (Range109-1862), HBV DNA (2650-118000000 IU/ml), sAg (250-125000 IU/ml) and with 19% eAg+ positivity. The demographic characteristics and clinical parameters at baseline for 16 rHBV, 7 nCHBV and 7 healthy controls are summarized in (Table S1).

### Epigenetic alterations in PBMCs of chronic Hepatitis B infected patients

The heatmap generated by unsupervised clustering of 549 significant genes revealed 284 hypomethylated and 265 hypermethylated genes (Fig.1A). Further, volcano plot revealed that epigenetic remodellers like KDM1B, KDM2B, JMJD1C, GATA6 and HDAC4 were hypermethylated in nCHBV (Fig.1B). These remodellers are known to play essential role in IL- 3 and IL-6 mediated signalling and increased CTLA4 expression. [25–27] Additionally, RPTOR, a metabolic gene and master regulator of metabolism along with COX19, CD247, CD300A, and IL4l1 were hypermethylated in nCHBV (Fig. 1B). Further, InnateDB and epigenes database identified125 differentially methylated immune genes and 32 epigenetic genes (Fig. S2A-B). Gene enrichment analysis by KEGG database revealed that immune associated genes like TOX, IL1RA, IL3RA, IL4R, IL17RB, IL-27, CD59 and CD99 were hypomethylated but CD247, RUNX3, CD300A, STAT5B, IL4l1, PRKCQ, ACVRL1 were hypermethylated. These genes in turn regulate Th1/Th2, Th17 cell differentiation, TGF-β signalling pathway, leukocyte trans-endothelial migration and cytokine receptor interaction (Fig.1C). mRNA expression analysis showed down regulation of TGF-β, KDM2B and up regulated expression of exhaustion marker PDCD1 in nCHBV (Fig. S2C).

**Fig. 1.**
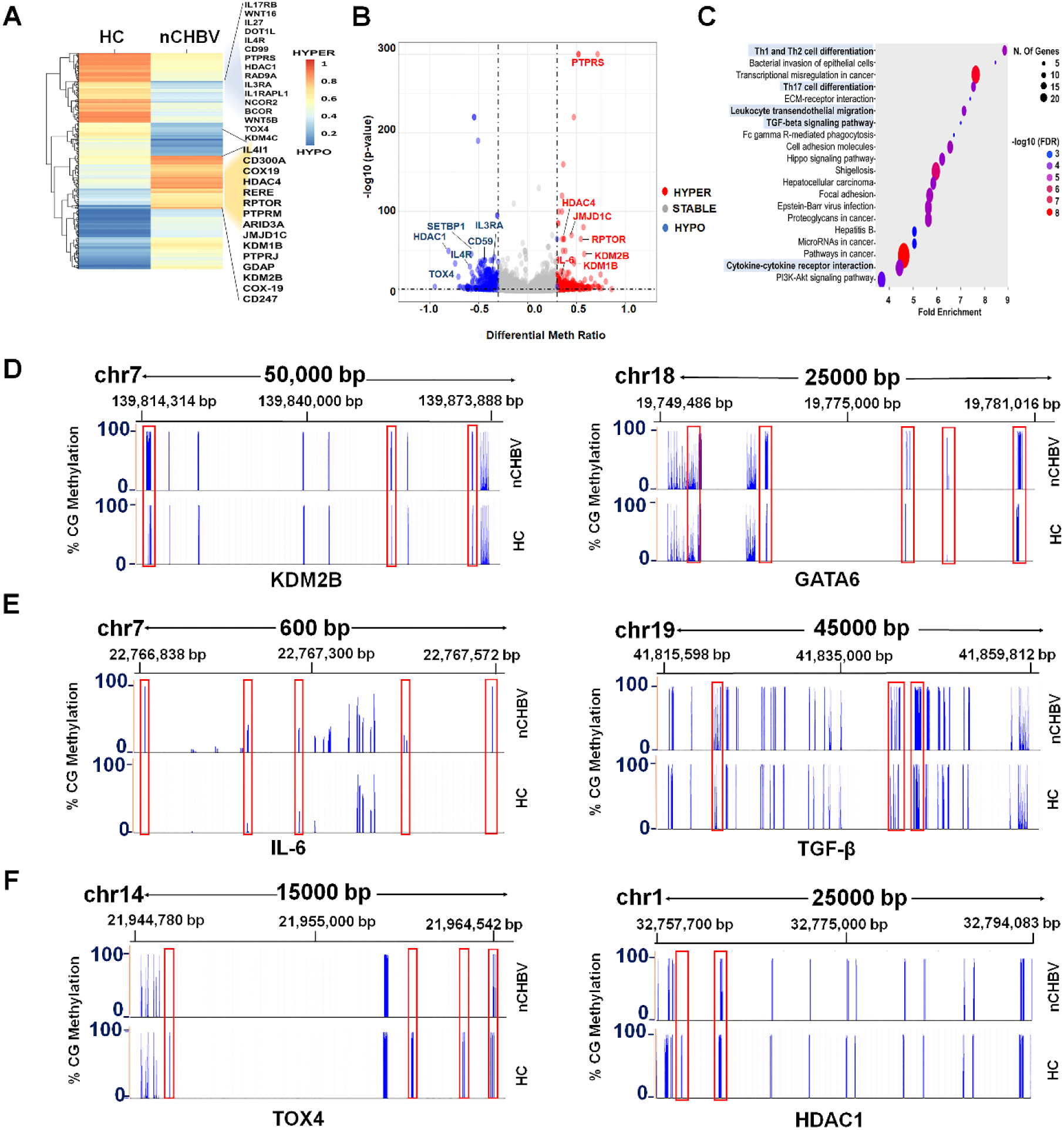
Genome wide methylation reveals novel epigenetic changes in nCHBV patients. (A) Heatmap showing unsupervised clustering of hyper and hypomethylated genes in HC and nCHBV (p=<0.05). (B) Volcano plots show differential meth ratio (>0.33) vs. log10 p value of selected genes in nCHBV patients compared to HC. (C) GSEA depicts genes enriched in nCHBV patients compared to HC. (D-E) Genomic view of KDM2B (D, Left panel), GATA6 (D, Right panel, IL6 (E, Left panel), TGF-β (E, Right panel) and) depicting hypermethylation at various locus in nCHBV. (F)Genomic view of TOX4 (Left panel), HDAC1 (Right panel), depicting hypomethylation at various locus in nCHBV compared to HC. HC, healthy control; nCHBV, naïve chronic hepatitis B infected.

Further, we have assessed the specific methylation positions in CpG islands of epigenetic remodellers KDM2B, GATA6, HDAC1 and immune genes, IL-6, TGF-β, TOX4. Our results showed that in KDM2B gene, 48 CpG sites were hypermethylated precisely in between position; 139814314 – 139873888 and 20 CpG islands of GATA6 between 19749486 -19781016 showed hypermethylation in nCHBV (Fig. 1D). As KDM2B regulates IL-6 [28], we have observed hypermethylation in 11 CpG islands of IL-6 promoter region at position; 22766838 -22767572 and 70 CpG sites of TGF-β at position; 41815598 - 41859812, indicating epigenetic remodellers mediated silencing of immune activating genes (Fig.1E). Indeed, TOX4 had hypomethylation of 7 CpG islands in promoter region at position 21944780 to 21964542 and 16 CpG islands of HDAC1 at 32757700 and 32794083 were hypomethylated in nCHBV implying a defect in downstream signalling leading to immune suppression (Fig.1F).

Further, while correlating with clinical parameters we have observed that hypermethylated epigenetic remodellers KDM1B, KDM2B, JMJD1C were negatively correlated with albumin, AST and HBVDNA respectively (Fig. S2D, left panel). However, HDAC4 showed positive correlation with total bilirubin while RPTOR shows positive correlation with AST. (Fig. S2D left panel, Table S2). Similarly, correlation analysis of hypomethylated genes including KDM4C, HADC1, IL17RB, showed negative correlation with anti-HBs and HBsAg (Fig. S2D, right panel). Both CD59 and IL3RA also depicted negative correlation with ALT (Fig. S2D, right panel).

### Distinct change in chromatin landscape in rHBV patients at the time of reactivation

From immune compromised state of chronic HBV to immune active state in reactivation, a sudden change in methylation landscape was evident in Principal component analysis (PCA) with clear separation between nCHBV and rHBV (Fig. 2A). Unsupervised clustering revealed 754 significantly differentially methylated genes during reactivation (Fig. 2B). Further, volcano plot defined epigenetic alteration with 640 hypo and 114 hyper-methylated genes in rHBV (Fig. 2C). It was observed that epigenetic remodellers, which were hypermethylated in nCHBV condition, showed prominent hypomethylation in rHBV. Further, we have observed epigenetic remodellers such as NCOR2, SETD7 and SIRT7 that were hypomethylated in healthy subjects, showed hypomethylation in rHBV (Table 1). In addition, immune genes IRF8, TGF-β1, IL12A, NFIX, NFIA, TCF21 and metabolic genes RPTOR, HK3, G6PD, GDPD1 and NADK were significantly hypomethylated in rHBV (Fig. S3A-B, Table 1). KEGG pathway enrichment showed Th17 differentiation, FOXO signalling, inflammatory mediated regulation of TRP channel, C-type lectin receptor signalling, leukocyte trans-endothelial signalling and hippo signalling pathways were enriched among the nCHBV and rHBV (Fig. 2D). Our flow cytometry analysis also depicted more IL-17, IFN-γ HBV specific CD8, Tfh cells as well as Th1/Th17 cell differentiation in rHBV compared to nCHBV (Fig. S3C).

**Fig. 2.**
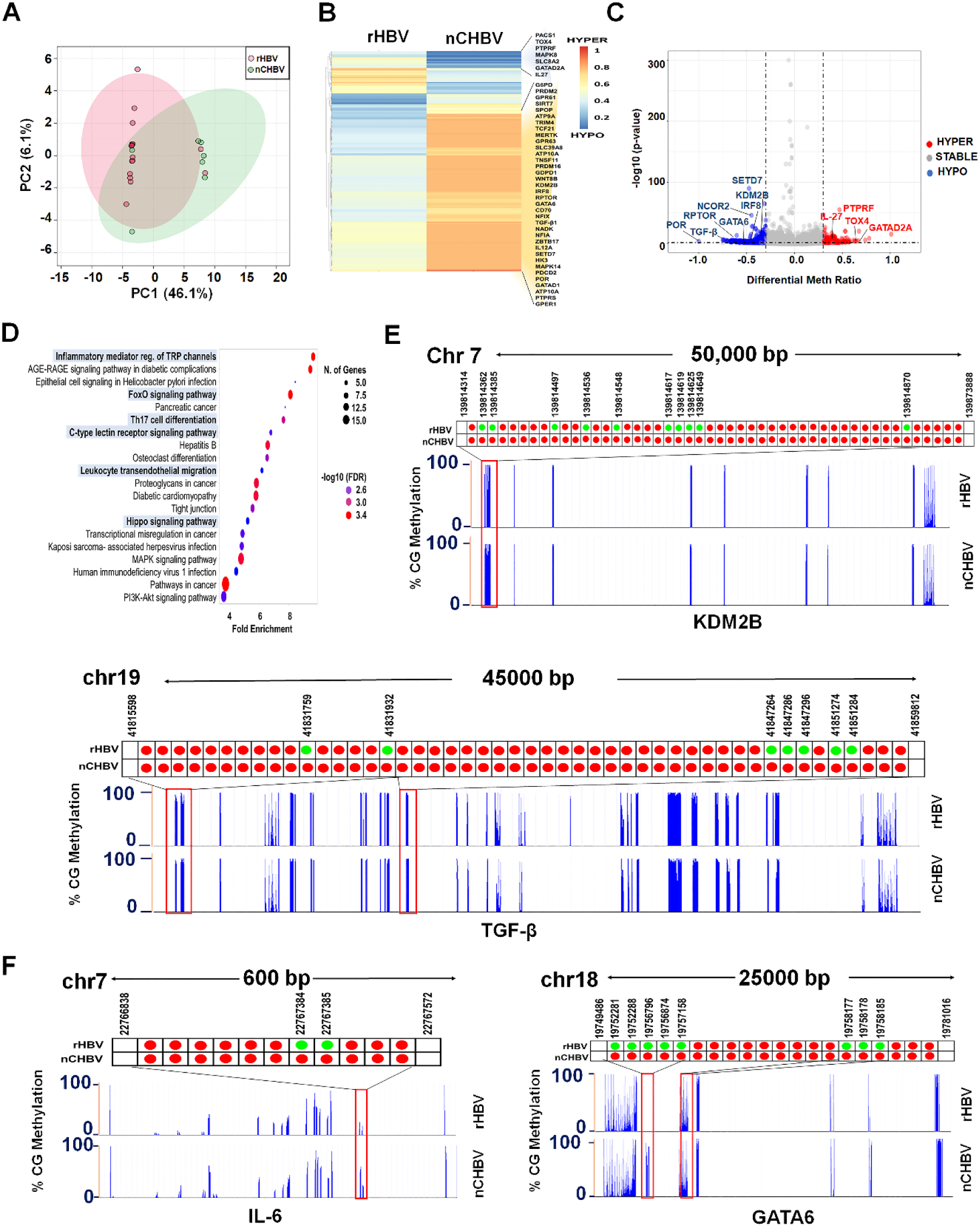
DNA methylation landscape of rHBV shows extreme hypomethylation of immune and epigenetic genes. (A) Principal Component Analysis of DNA methylation in rHBV compared to nCHBV. (B) Hierarchical clustering representation of 793 genes identified as differentially methylated in rHBV and nCHBV. (C) Volcano plots show meth ratio (>0.33) vs. -log10 p value (p<0.05) of selected genes in rHBV patients compared to nCHBV. (D) GSEA depicts KEGG database pathway enrichment in rHBV patients compared to nCHBV. (E-F). Genomic view of genes depicting hypomethylation at various locus in rHBV compared to nCHBV; KDM2B (E, Upper panel), TGF-β (E, Lower panel), IL6 (F, Left panel) and GATA6 (F, Right panel). nCHBV, naïve chronic hepatitis B infected; rHBV, HBV reactivation.

**Table 1:**
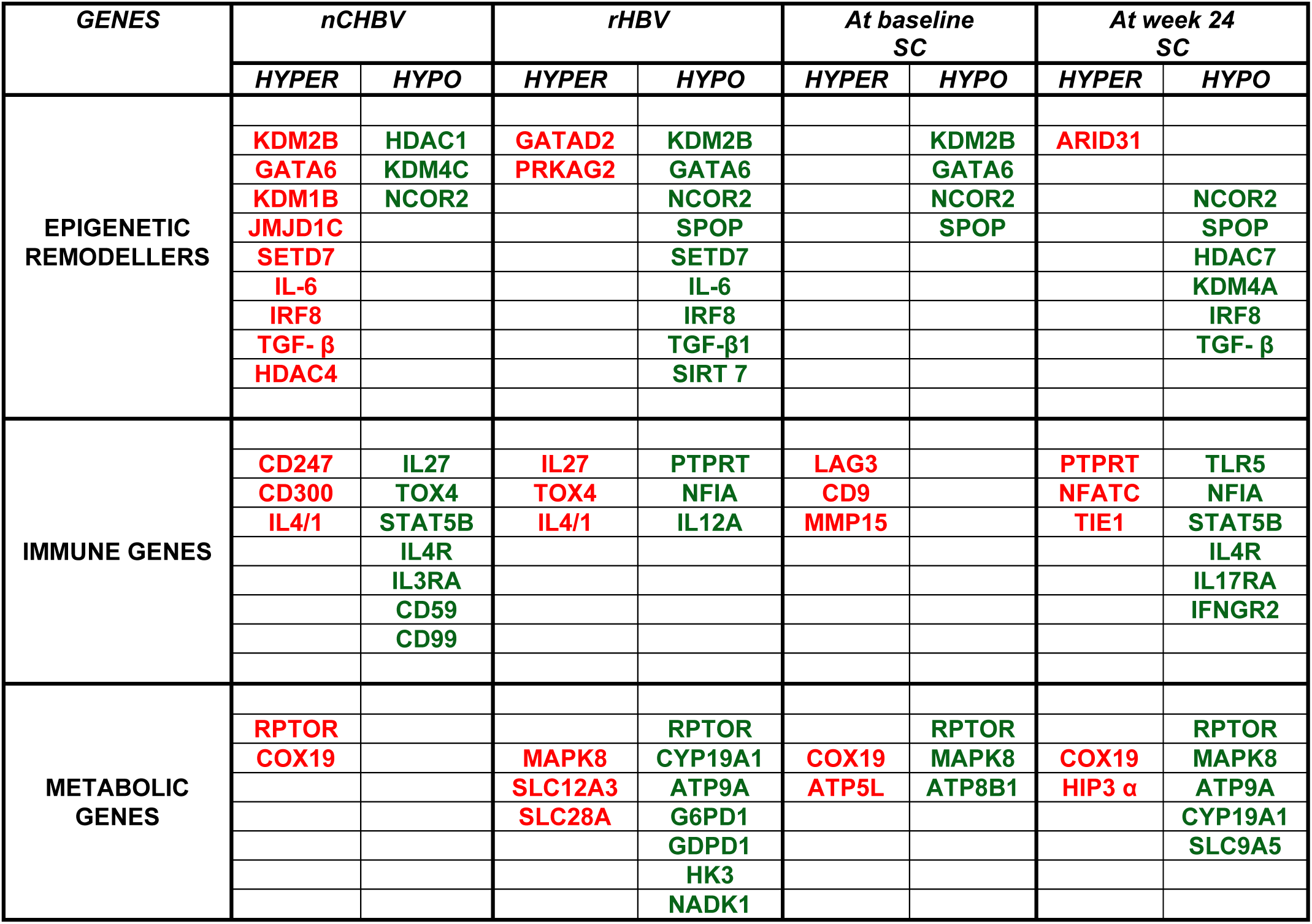
Hyper and hypomethylated genes in nCHBV, rHBV and Seroconveters.

Interestingly, we have found that out of 48 hypermethylated CpG islands of KDM2B in nCHBV, 10 CpG islands were hypomethylated in rHBV at specific positions (Fig. 2E). Similarly, 7 CpG islands of TGF-β, acquired hypomethylation at positions 41815598 to 41859812 (Fig. 2E). Further, out of 11 hypermethylated CpG islands of IL-6 gene, 2 CpG islands gained hypomethylation at 22767384 and 22767385 positions in rHBV (Fig. 2F) and 6 CpG islands of GATA6 depicted hypomethylation in reactivation as compared to nCHBV (Fig. 2F). By using TRANSFAC 4.0, we have assessed binding affinity for transcription factors in the hypomethylated CpG islands of IL-6 for various transcription factors like GATA1, TBP, OCT-1 and NF-1 (Fig. 3A). Similarly, hypomethylated TGF-β CpG islands also showed affinity for CEBPα, Sp1, OCT-1 and SOX9 transcription factors (Fig. 3B).

**Fig. 3.**
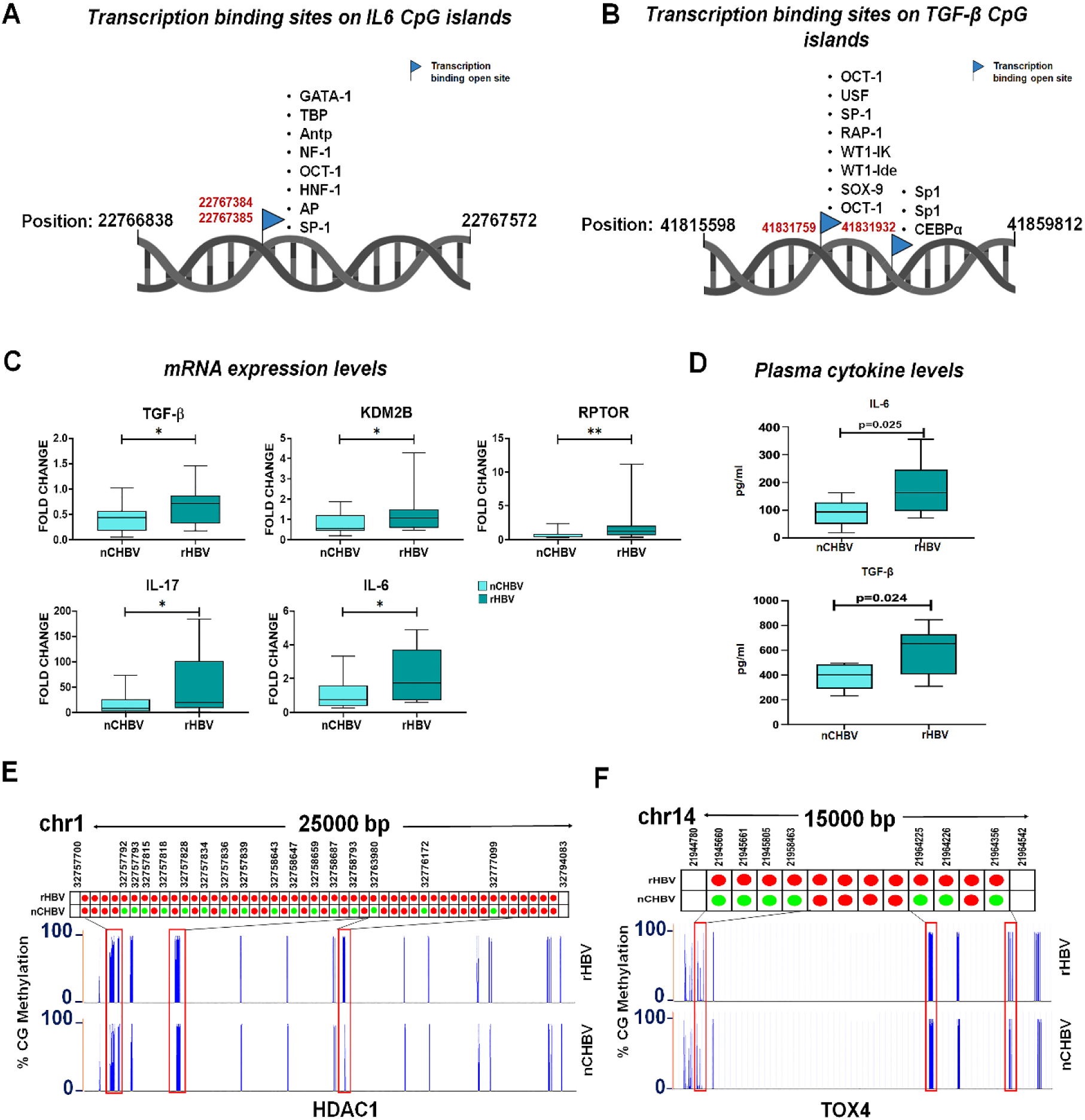
DNA hypomethylation of epigenetic and immune genes at the time of reactivation. (A-B) Binding sites for various transcriptional factors (TF) at hypomethylated region of IL-6 and TGF-β. (C) mRNA expression levels of TGF-β, IL- 17, KDM2B, IL6 and RPTOR in rHBV and CHBV. (D) Plasma cytokine levels of IL-6 and TGF-β. (E-F) Genomic view of HDAC1 and TOX4 indicating hypomethylation at promoter locus in rHBV compared to nCHBV. nCHBV, naïve chronic hepatitis B infected; rHBV, HBV reactivation.

At the same time, mRNA expression analysis revealed increased expression of KDM2B, IL-6, TGF-β, IL-17 and RPTOR in rHBV (Fig. 3C). Additionally, this data was also supported by the plasma concentrations of significantly increased IL-6 and TGF- β levels in rHBV (Fig. 3D). Further, hypomethylated KDM2B gene was positively correlated with clinical parameters like PT, INR, MELD and MELD-Na while negatively correlated with albumin (Fig. S3D, Table S3). Correlation analysis also showed positive correlation of PRDM16 and SETD7 with albumin while GATA6 was negatively correlated (Fig. S3D).

However, HDAC1 and TOX4 genes, which were hypomethylated in nCHBV, showed hypermethylation status in rHBV. 50 CpG islands of HDAC1 from 32757700 to 32794083 positions showed hypermethylation whereas 11 CpG islands of TOX4 from 21944780 to 21964542 positions were hypermethylated in rHBV (Fig. 3E-F).

TOX is a master regulator of all inhibitory molecules and our flow cytometry data revealed downregulated expression of TOX on HBV specific CD4+ and CD8+ T cells in rHBV compared to nCHBV (Fig. S3E-F). In summation, these data indicate that epigenetic remodellers mediate immune and metabolic reprogramming during HBV reactivation.

### HBsAg seroconverters showed hypomethylation of IRF8 and TLR5 at baseline

There were no significant difference in demographic and clinical parameters in SC and NSC at baseline (Table S4), however, their epigenetic profile was markedly different in SC patients. Principal component analysis depicted a significant difference in methylation status of SC and NSC (Fig. S4A). Unsupervised clustering of 233 significant genes (161 hypomethylated and 62 hypermethylated) were associated with WNT, mTOR and Hippo signalling, inflammatory mediator regulation of TRP channel, and chemokine pathways (Fig. S4B-C). HBsAg seroconversion is a functional cure for chronic HBV infection, therefore by analysing position specific methylation pattern we tried to delineate the genes which can predict seroconversion at baseline. At baseline, in SCs, LAG3, CD9 and MMP15 were hypermethylated while IRF8, TLR5, TGF-β, and GATA6 were hypomethylated (Fig. S4D). At the same time, flow cytometry data revealed that expression levels of TOX on HBV specific CD4 and CD8 T cells were significantly decreased (p<0.05) in SC compared to NSC (Fig. S4E: Left and Right panels).

In addition, metabolic reprogramming in SC depicted hypomethylation of RPTOR, ATP8B1, COX19, SLC25A19 and SLC26A16 genes (Fig. S4D). Furthermore, genes such as NCOR2 and SPOP, which play an important role in immune activation and differentiation were also hypomethylated in SC compared to NSC (Fig. S4F).

We have also analysed methylation islands of LAG3. It was observed that 14 CpG sites were hypermethylated from position 6881700 to 6887544 in SC but hypomethylated in NSC. This could be the cause of immune exhaustion in NSC patients (Fig. 4A). However, at baseline, interferon related IRF8 was hypomethylated at 7 CpG islands from 85932416 to 85954818 in SC compared to NSC (Fig. 4B), which depicted that Jak-Stat signalling and cytokine secretion. Additionally, TLR5 also depicted 4 hypomethylated CpG sites at position 223286640 to 223316610 in SC than NSC (Fig. 4C). mRNA expression and plasma cytokine levels in SC had increase in KDM2B, TGF-β and IL-6 expression levels (p<0.05), which is suggestive of immune differentiation (Fig. S4D). Hypomethylation of IRF8 and TLR5 CpG islands depicts open sites for various transcription factor like Sp1, GATA1, IRF1, ISGF-3 (Fig. 4D) and NF1, OCT-1 respectively (Fig. 4E) leading to significantly higher mRNA expression of IRF8. COX19 also showed similar expression (Fig. 4F).

**Fig. 4.**
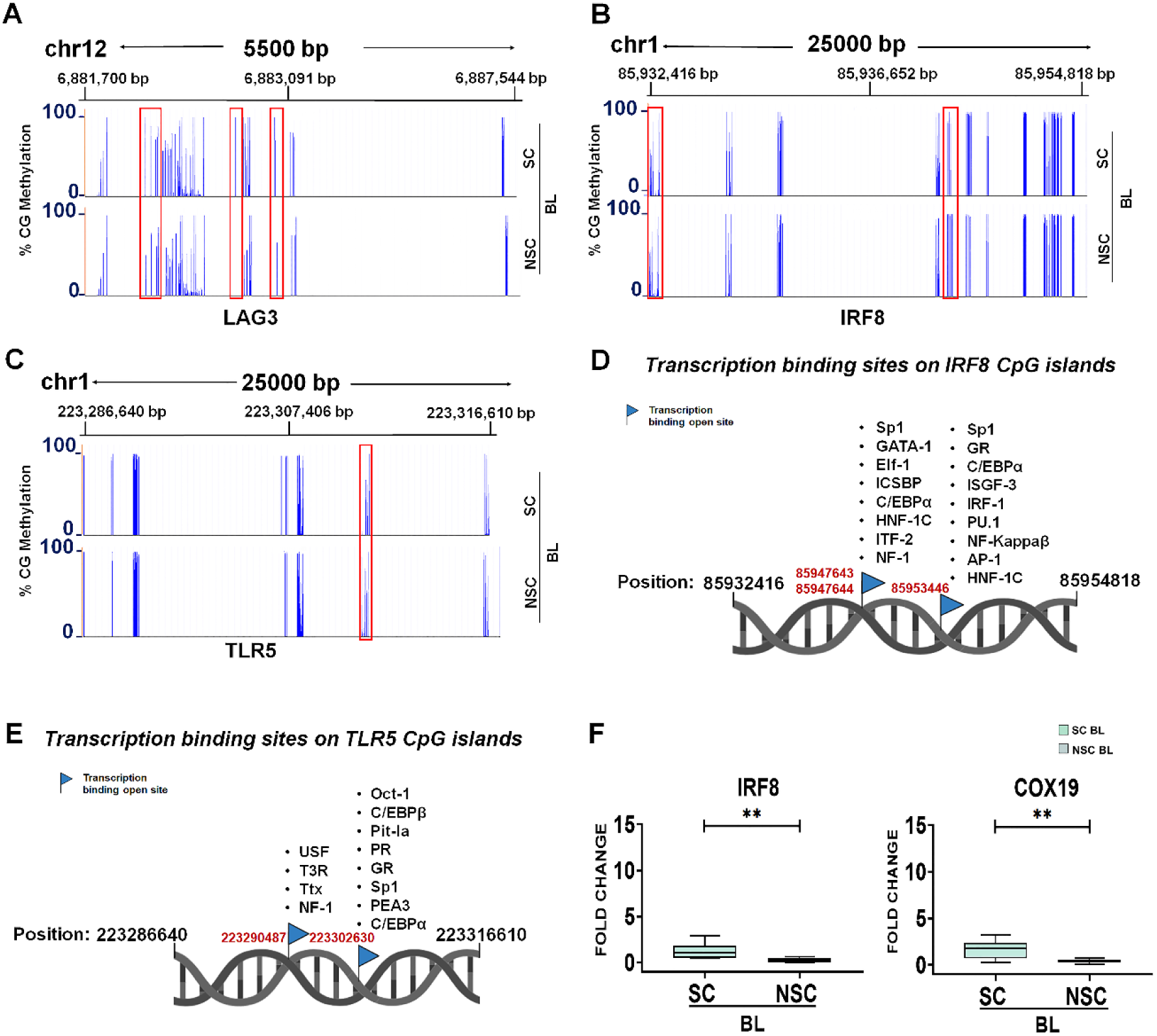
At baseline, distinct pattern of methylation in seroconverters and non-seroconverters. (A) Genomic view of LAG3 indicating hypermethylation at various positions in SC compared to NSC. (B-C) Genomic view of hypomethylated IRF8 and TLR5 in SC compared to NSC. (D-E) Transcription factors binding site in hypomethylated region of IRF8 and TLR5 gene. (F) mRNA expression levels of IRF8 and COX19 in SC and NSC. SC, Seroconverters; NSC, Non- seroconvertors.

This data suggests that increase in KDM2B expression leads to increase in cytokine production such as IL-6 and TGF-β, which further increase Th1/17 cells differentiation. In conclusion, hypermethylation of LAG3 and hypomethylation IRF8 and TLR5 distinctly segregate SC patients from NSC at baseline.

### Hypomethylation and Chromatin accessibility of IL17RA, IFNGR2 and TLR5 in seroconverters at week 24

At week 24, in SC, epigenetic landscape showed further changes. Along with IRF8 and TLR5, other immune, metabolic and epigenetic genes were hypomethylated including IL17RA, IFNGR2, KDM4A, HDAC7, NCOR2, TGF-β, NFIA, STAT5B, MAPK8, CYP19A1, ATP9A and SLC9A5 (Fig. 5A-B, Fig. S5A-B). KEGG pathway depicted hypomethylated genes associated with Th17 cells differentiation, Wnt signalling, Th1 and Th2 cells differentiation, leukocytes trans-endothelial migration and Hippo signalling (Fig. 5C). Further, site specific methylation analysis revealed that in IL17RA, 12 CpG islands were hypomethylated at position 17565868 to 17594892 (Fig. 5D) in SC compared to NSC while IFNGR2 showed significant hypomethylation of 5 CpG from 34775451 to 34851145 positions at week 24 (Fig. 5D). TLR5 had 10 CpG loci hypomethylated starting from position 223286640 to 223316610. Transcription factors TBP, OCT1, Sp1 and NF1 showed increase binding affinity to IFNGR2 at hypomethylated (open) sites (Fig. 5F). Open transcription factor binding sites for IL17RA were also analysed (Fig. S5D). mRNA expression analysis showed increase in IL17, TGF-β, IRF8, and COX19 in seroconverters (Fig. S5E). Similarly our flow Cytometry data also suggested significant increased IL-17, IFN-γ secreting HBV specific CD4 and CD8 T cells in SC compared to NSC (Fig. S5F).

**Fig. 5.**
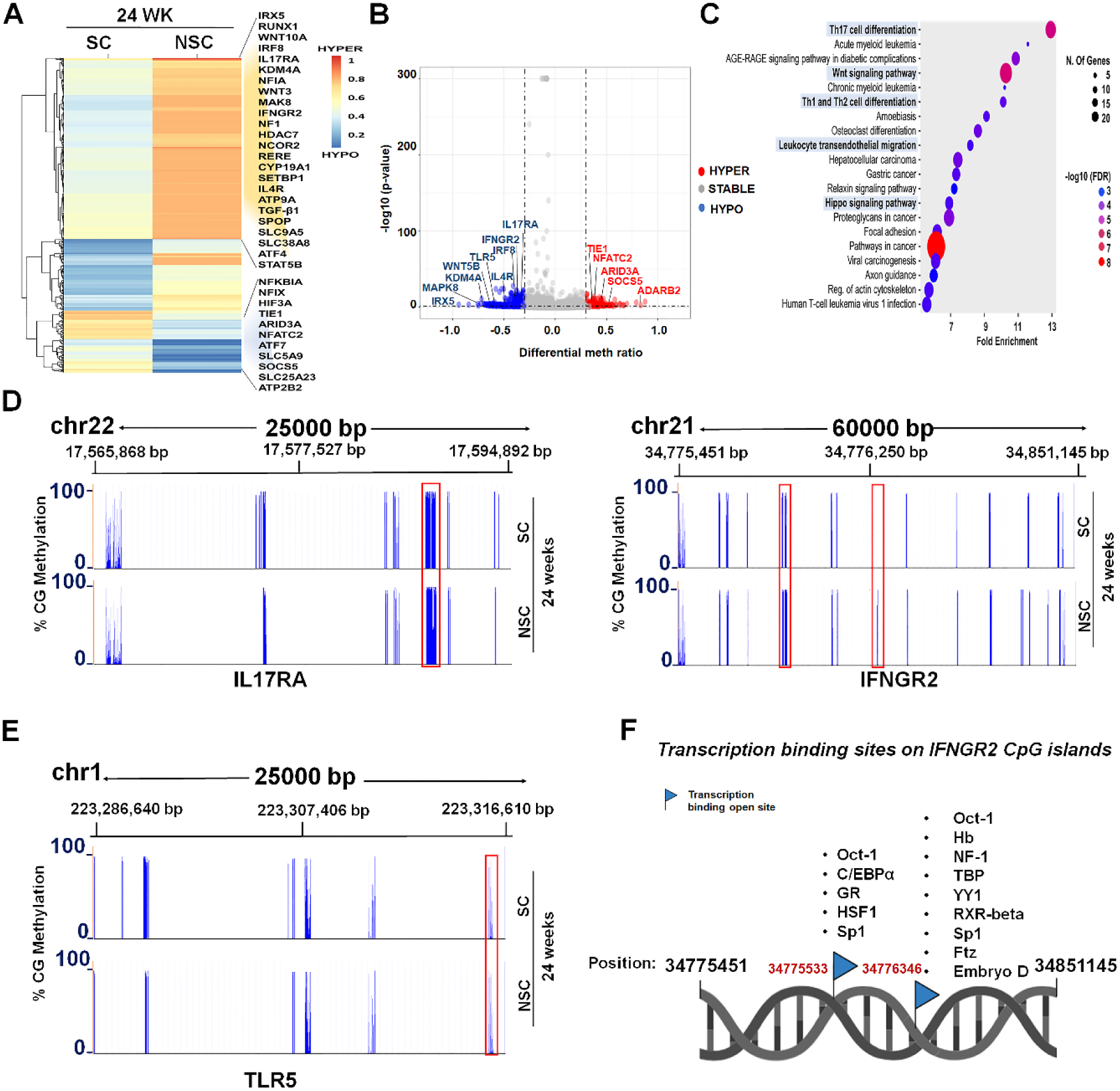
Epigenetic changes at 24 weeks in seroconverters and non-seroconverters. (A) Differentially hyper and hypomethylated genes in SC and NSC. (B) Volcano plots show meth ratio (>0.33) vs. -log10 p value (p<0.05) of selected genes in SC compared to NSC. (C) GSEA depicts pathway enrichment in SC compared to NSC. (D) Genomic view of IL17RA (left panel) and IFNGR2 (right panel) indicating hypomethylation at various locus in SC compared to NSC. (E) Genomic view of human TLR5 indicating hypomethylation at various locus in SC compared to NSC. (F) Affinity for binding of transcription factors in hypomethylated region of IFNGR2.

## Discussion

HBV persistence is due to inability of the immune system to resolve chronic hepatitis B infection. [29] Till now, two major forms of epigenetic regulation are known in HBV induced HCC: posttranslational modification of histone proteins associated with the cccDNA mini chromosome and DNA methylation of viral and host genomes. [30] HBc, HBx and cccDNA directly interact with several host factors like p300, KDM1A, CREB1, HDAC1, PRTM5, E2F1 and SIRT1 to modulate their expression for immune escape. [30–31] Therefore, epigenetic factors contribute to the outcome of chronic hepatitis B virus (HBV) infection by affecting immunity as well as viral replication. We have observed the altered methylation status of epigenetic remodellers like KDMs, JMJD1C, GATA6, HDACs and immune genes like IL6, IL17, TGF-β, CD300A, and IL4 in nCHBV. Further, immune exhaustion genes like TOX and CD59 were also hypomethylated in nCHBV. TOX is a master regulator of PDCD1, LAG3, TIM3 and other exhaustion genes and TOX hypomethylation leads to increased expression of these molecules. [32–33] Increased expression of PD1 in CD8+ T cells in chronic HIV patients also demonstrated sustained unmethylated PD-1 locus after prolonged exposure of HIV virus. [34] mRNA expression of PDCD1 and decreased plasma levels of IL-6 and TGF-β is suggestive of immune exhaustion in nCHBV (Fig.1 and Fig.S2).

HBV reactivation is the reappearance of necro-inflammatory liver disease in CHBV patients due to immunosuppression or stopping of antivirals. It involves reprograming of immune, metabolic and epigenetic genes.[35] Therefore, we observed changes in methylation status of epigenetic, immune and metabolic genes during HBV reactivation which plays an important role in the course of seroconversion. Epigenetic modulation influences disease patterns. Recently, it was observed that low chromatin accessibility of TBX21, NFκB1, TP53, STAT1, MAFK and RUNX3 genes in CD4+ and CD8+ T cells contribute to pathogenesis in moderate to severe COVID patients. [36]

Histone modification and DNA methylation plays an important role in differentiation of naïve cells into mature T cells. It was observed that specific epigenetic modifiers regulate T cell differentiation. [37] Upon antigen recognition, key effector genes such as Gzmb and Prf1, acquire hypomethylation to gain chromatin accessibility in mature T cells, while naïve-associated genes remain repressed. [38] Also, for pathogen clearance, memory precursor cells differentiate into long-lived memory cells by demethylating IL17RA and Bcl2. [38] We have also observed altered methylation status with enrichment of Th17 differentiation and hippo signalling in rHBV. Our immune profiling data of T cells also reveal increased frequencies of IL-17, IFN-γ secreting HBV specific CD8, Tfh as well as Th1/17 cells in reactivation compared to nCHBV. Indeed, Th1/17 cells differentiation was supported with hypomethylation status of many epigenetic remodellers like KDM2B, NCOR2, PRDM16, GATA6 and SETD7 driving IL-6, TGF-β signalling and immune reprogramming by increasing type I interferon response, immune activation, B cell differentiation and trained immunity.[39–42] Another molecule SETD7, known as metabolic gatekeeper of mTOR signalling was hypomethylated in reactivation suggesting the reprogramming of metabolic pathway.[43] Chronic HBV patients show dysregulation of TCA cycle and shift towards lactate production leading to T cell exhaustion.[44–45] In rHBV patients, metabolic genes mainly RPTOR, G6PD1, GDPD1 and NADK were hypomethylated which regulates TCA cycle, oxidative phosphorylation, ATP production and mTOR signalling.[46] In addition, hypomethylation of IL-6, TGF-β, KDM2B, GATA6 and TCF21 showed binding affinity for transcription factors like TBP, OCT-1, SP1 and NF-1 in rHBV. It is evident from previous studies, OCT-1 has important role in T-cells development and differentiation (mainly Th1 and Th17).[47] TBP binds to promoter site of various genes and regulates their expression. Gene SP1 transcriptionally activate NLRP6 inflammasome. [48] TCF21 promotes pro-inflammatory cytokine IL-6 production and also regulate matrix metalloproteins (MMPs) expression. [49]

In reactivation, there is a resurge of virus with an abrupt alteration in liver functions with more chances of coagulopathy, ascites and hepatic encephalopathy. However, gene correlation with clinical parameters was also observed. Hypomethylated IRF8 showed positive correlation with increased albumin production, indicating better liver functionality. This finding strengthens the notion of epigenetic mediated immune and metabolic reprogramming during reactivation.

In reactivation, there is a probability of HBsAg clearance by raising anti-HBs titres (>10mIU/ml), known as seroconversion. Therefore, we tried to delineate seroconverters and non-seroconverters at the time of reactivation (baseline) by analysing methylation status. Our study nicely proved distinct methylation patterns in SC and NSC at baseline. In SC, KDM2B, GATA6 and NCOR2 epigenetic remodellers were hypomethylated at baseline. Immune genes IRF8, TLR5 and COX19 were also hypomethylated especially position specific hypomethylation of IRF8 and TLR5 in SC. These hypomethylated sites were opened for binding of GATA1, SP1, NF-κB, NF-1 and CEBPα transcription factors. Gene expression results also showed increased expression of IRF8 and COX19. Similarly, LAG3 showed hypermethylation at various positions supported by significantly decreased mRNA expression of TOX and PD1 exhaustion genes in SC. This decreased TOX expression in SC is also supported by our flow cytometry data. Therefore, at baseline, there was reprogramming of immune genes in SC as compared to NSC.

At week 24, IL17RA, IFN-γ, TGF-β, NFIA, STAT5B and CD70 were hypomethylated along with IRF8, COX19, TLR5 and GATA6 in SC. Position specific hypomethylation of IFNGR2, IL17RA, IRF8 and COX19 showed affinity for binding of OCT-1, Sp-1, NF- 1, CEBPα and CEBPβ. A significant increase in TGF-β, IRF8, CD70 and GATA6 plays crucial part in Th17 cell differentiation, Jak-STAT signalling, loss of exhaustion markers and Tregs. Indeed, in SC, increased TGF-β hypomethylation was positively correlated with increase expression of anti-HBs titres.

Therefore, it can be concluded that hypomethylation of epigenetic, immune and metabolic genes and hypermethylation of exhaustion genes at baseline and week 24 suggest immune metabolic reprograming in SC but defective in NSC.

## Limitations of the study

This study has following limitations: We had small cohort of chronic HBV infected patients, which were randomly recruited. We have also showed transcription binding affinity on open sites using online tool TRANSFAC 4.0 but we did not investigate direct impact of these transcription factors. This study does not include any inhibitors of epigenetic remodellers and their effect on immune resetting.

However, specific epigenetic signatures in seroconversion are our novel finding. This may be use therapeutically for the development of drugs to cure chronic HBV infection and non-seroconverters in HBV reactivation.

## Author’s contributions

JKS and MI: Performed DNA methylation experiments, data analysis, interpretation, and initial draft of manuscript writing. GV, AS, PB and SP: Helped in initial data analysis. GR: Provided critical inputs for data analysis, reviewed analysis. AJ, MKS and SKS: Patient stratification, clinical data review. SKS: Critical inputs in manuscript editing. NT: Conceptual design of study, designing of experiments and analysis, critical revision of the article for important intellectual content and finalised the manuscript.

## Acknowledgement

We would like to thank all the patients who participated in the study and Dr. Surabhi Srivastava, from Centre for Cellular and Molecular biology (CCMB), Hyderabad for initial input in methylation data analysis.

