## Supplementary information for "Epigenetic mediated functional reprogramming of immune cells leads to HBsAg seroconversion in Hepatitis B Virus Reactivation patients"

**Methods**

**Retrospective Study Cohort:** In our previous study, we have collected 67 reactivation patients, nineteen naïve CHBV and ten healthy subjects (Islam et al. 2022) .^[35]^. Within 24-48 weeks, 12 rHBV patients had loss of HBsAg and developed anti-HBS titres (>10 IU/mL), which are considered as seroconverters (SC) and rest as non-seroconverters (NSC) (Study Cohort Fig. S1). Most of the seroconverters develop anti-HBs titres within 24 weeks. Patients having co-infections with Hepatitis A, C, D, E or HIV, or other causes of chronic liver failure, hepatocellular carcinoma (HCC), portal vein thrombosis, coexistent renal impairment and pregnancy were excluded from the study.

Healthy subjects were age and sex matched with no prior history of any liver related diseases. They must be negative for all hepatitis viral markers: Immunoglobulin M (IgM), Hepatitis A virus (anti-HAV), Hepatitis E virus (anti-HEV), Hepatitis B surface antigen (HBsAg), anti-HCV, HIV, CMV and HSV.

This study protocol was approved by the ethics and review committee of the Institute of Liver and Biliary Sciences (ILBS) (No. IEC/2017/49/ NA02) and was in accordance to the ethical guidelines of the declaration of Helsinki. All participants provided a written Informed consent.

10-12 ml of peripheral blood was collected from all patients and healthy subjects in EDTA tubes. Plasma was separated and serological and virological markers including HBsAg, anti-HBs, anti-HBc, HBeAg, anti-HBe, HBV DNA, cytokine bead array were measured as detailed in Islam et al. 2022.^[35]^. T cell functionality was studied using Flow cytometry and data was reanalysed for our present study cohort which were used for methylation studies.^[35]^

**Reduced Representation Bisulfite library construction and sequencing:**

8-10 ml peripheral blood was processed for PBMCs isolation. From PBMCs, we have isolated DNA for preparing Reduced Representation Bisulfite Sequencing (RRBS) libraries using Zymo-Seq RRBS Library Kit (Zymo Research, USA, #D5461). In brief, 300ng of genomic DNA was purified with a GeneJET Genomic DNA purification kit (Thermofisher, USA, #K0721) and digested with restriction enzyme MspI (5’C’CGG) by incubating at 37°C for 4 hrs (Details in Supplementary data). ^[20]^

It is already known that CpG island are enriched with CCGG site at promoter and genic region of the genome. Msp1 is a restriction endonuclease which recognises at restriction site CCGG and cut between two Cs. However, this enzyme cuts all CCGG sites irrespective of the methylation status of Cs. RRBS generally captures approximately 80% of CpG islands and 60% of the promoter regions. Further, the CpG rich genomic fragments were ligated with linker adapters and gap was filled with 5-Methylcytosine dNTP mix. The ligated fragments were purified with Zymo spin- IC column. Later on, the DNA fragments were treated with lighting conversion reagent, which can chemically modify the non-methylated cytosine into uracil upon the addition with L- desulphonation buffer. Furthermore, the index primer amplification was done using standard dNTPs and a uracil-tolerant Taq DNA polymerase which generates libraries with thymine in place of the originally unmethylated cytosines. The amplified product was then purified using Zymo spin- IC column and the final library was eluted in 20µl of DNA elution buffer. Final product was sequenced and analysed.

**RRBS Sequencing Data processing and Analysis:**

RRBS analysis of rHBV, nCHBV and HC FastQC (version 0.11.8, www.bioinformatics.babraham.ac.uk/projects/fastqc/) was used for quality check. Universal adapter was removed using trim galore (version
0.6.2, https://www.bioinformatics.babraham.ac.uk/projects/trim_galore/), a wrapper script to automate quality and adaptor trimming as well as quality control. Genome indexing was done using Batmeth2. These reads were aligned to the reference Hg19 genome downloaded from Ensemble release 102. Mapping and alignment were performed using Batmeth2 using align tool. Also, DNA methylation level was calculated using Batmeth2 calmeth tool. Then annotation was done using MethyGff tool using hg19 genome gff and genome methylation obtained using calmeth tool. Methdiff.py from BSMAPZ tool (fork of BSMAP V2.90) was used for differential methylation ratio (DMRs) analysis. Selected gene methylations were fetched using perl and shell script.^[21]^ We have identified the immune genes using an integrated analysis platform, innateDB database. Similarly for epigenetic genes we used publicly available epigene database. Further, we have assessed Transcription factor binding sites by using TRANSFAC 4.0 software and the AliBaba 2.1 database.

**Methylation calculation and Analysis:**

Methylation ratio was calculated by Batmeth2 calmeth tool. Batmeth2 tool comprises different methods to detect differentially methylated regions and one of them is beta binomial distribution model for data with replicates. One of the major goals in methylation data analysis is to identify differentially methylated cytosines (DMCs) as well as differentially methylated regions (DMRs). Since, it is known that SNP variation also occurs from C to T, which may affect methylation levels in cytosine loci leading to calculation error. Therefore, to confirm C to T bisulfite conversion, it is necessary to simultaneously examine reverse complementary strand as well. The β value ranges from 0 to 1, where 0 indicates completely unmethylated and 1 indicates completely methylated sites. Meth ratio >±0.3 is considered as hyper and hypomethylation. ^[22-24]^ Methylation data represented in heatmap was done by “Pheatmap” tool. Data depicted in volcano plot were obtained using “Enhancedvolcano” tools. Correlation was done using “corplot” tools with R package 4.1.0.

**RNA isolation and quantitative-PCR (Q-PCR) assays:**

Total RNA was extracted from PBMCs using TRIzol reagent (Thermo Fisher) according to the instructions. Complementary DNA (cDNA) was synthesized using High capacity cDNA kit (ABI, Thermo Fisher Scientific, Waltham, MA, USA; #4368813) and qRT-PCR assays were performed for genes using following primers and SYBR Green Master Mix (Thermo Fisher; #A25742) in Quant Studio real time PCR machine (Thermo fisher). Data were normalized to GAPDH expression.

| **Gene name** | **Forward primer (5’-3’)** | **Reverse primer (5’-3’)** |
| --- | --- | --- |
| PDCD1 | CGCTTCCGTGTCACACAACT | AGGTAGGTGCCGCTGTCATT |
| IL-17 | ACGAAATCCAGGATGCCCAA | TCCGGTTATGGATGTTCAGGTT |
| IFN-γ | ACTGTCGCCAGCAGCTAAAA | TATTGCAGGCAGGACAACCA |
| TGF β | GAGGCGCCCGGGTTATGCTGGTTG | CGCAAGGACCTCGGCTGGAAGTGG |
| GAPDH | CCATCACCATCTTCCAGGAGCGA | TGCAAATGAGCCCCAGCCTTCTC |
| LAG3 | TGACTGGAGACAATGGCGAC | GGATTTGGGAGTCACTGTGATG |
| TOX | CCACCATGCAGCAAGGATTTA | CCCGAACGCACATACTCCAT |
| KDM2B | TGTAGATCTCGGCTCTGGCT | GTCAATCGGGCGCAATCTTC |
| IL-6 | CACTGGTCTTTTGGAGTTTGAGG | CACAGCTCTGGCTTGTTCCT |
| IRF8 | CGAGGTTACGCTGTGCTTTG | GCCACGCCTAGTTTGCATTTT |
| COX19 | AATGCAGGATGGAGAGAAAATTGA | CAGCCAACCTAAGAGGCAGG |
| RPTOR | AGCATCGGATGCTTAGGAGTGG | CAGCCAGTCATCTTTGGAGACC |

**Statistical Analysis:**

The comparison of continuous data was performed using one way ANOVA / Kruskal-Walli’s test and data are represented as mean± standard error of mean (SEM). Statistical analysis for flow cytometry, cytokine bead array and QRT- PCR data was performed using Prism (Graph Pad version 9.0). A p value <0.05 was considered significant. A student t test (non-parametric) Mann Whitney was used for comparison between two independent conditions. Pearson correlation analysis was performed using R package 4.1.0.

**Supplementary Figures:**

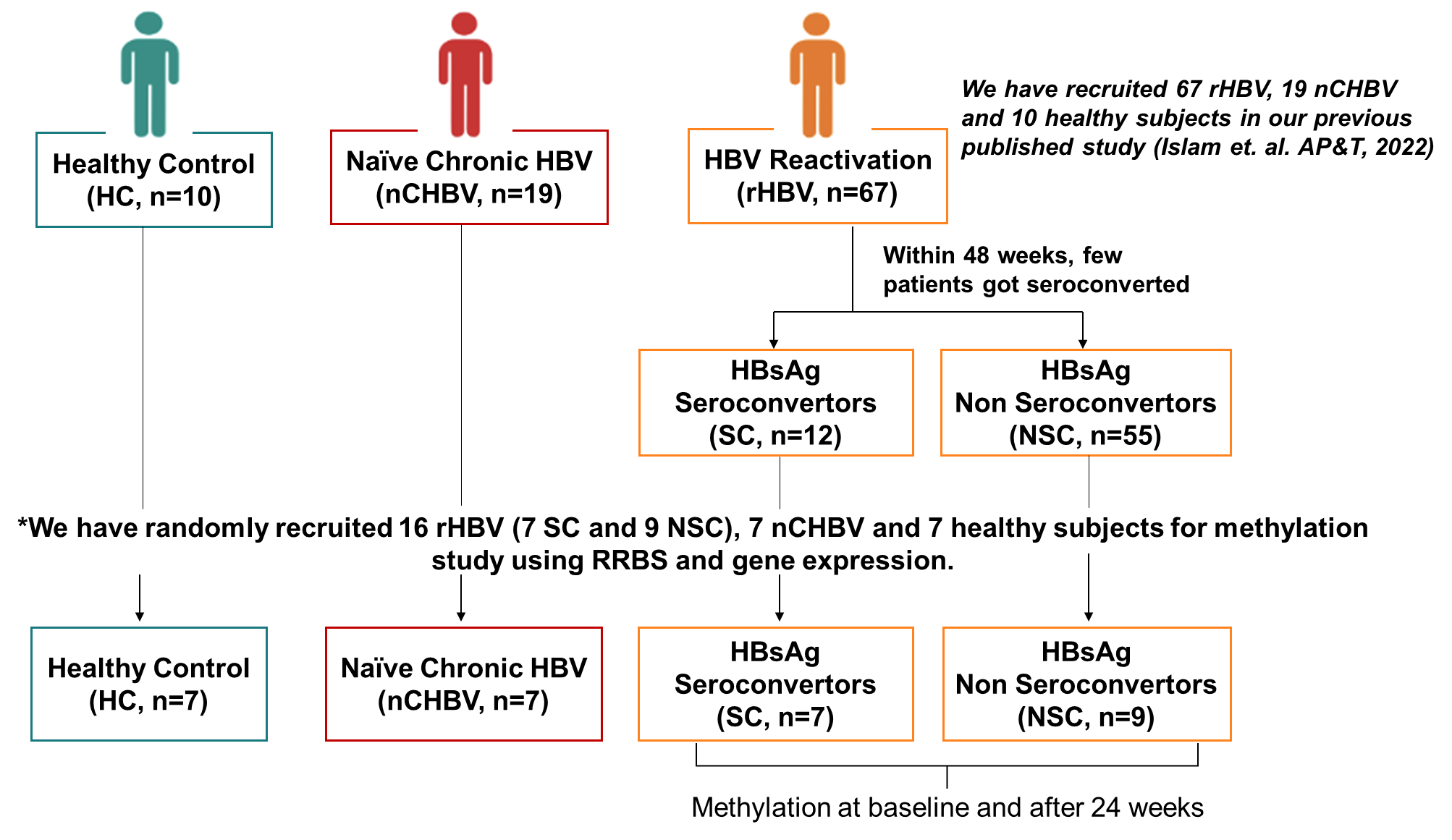

**Fig. S1. Consort Diagram.**

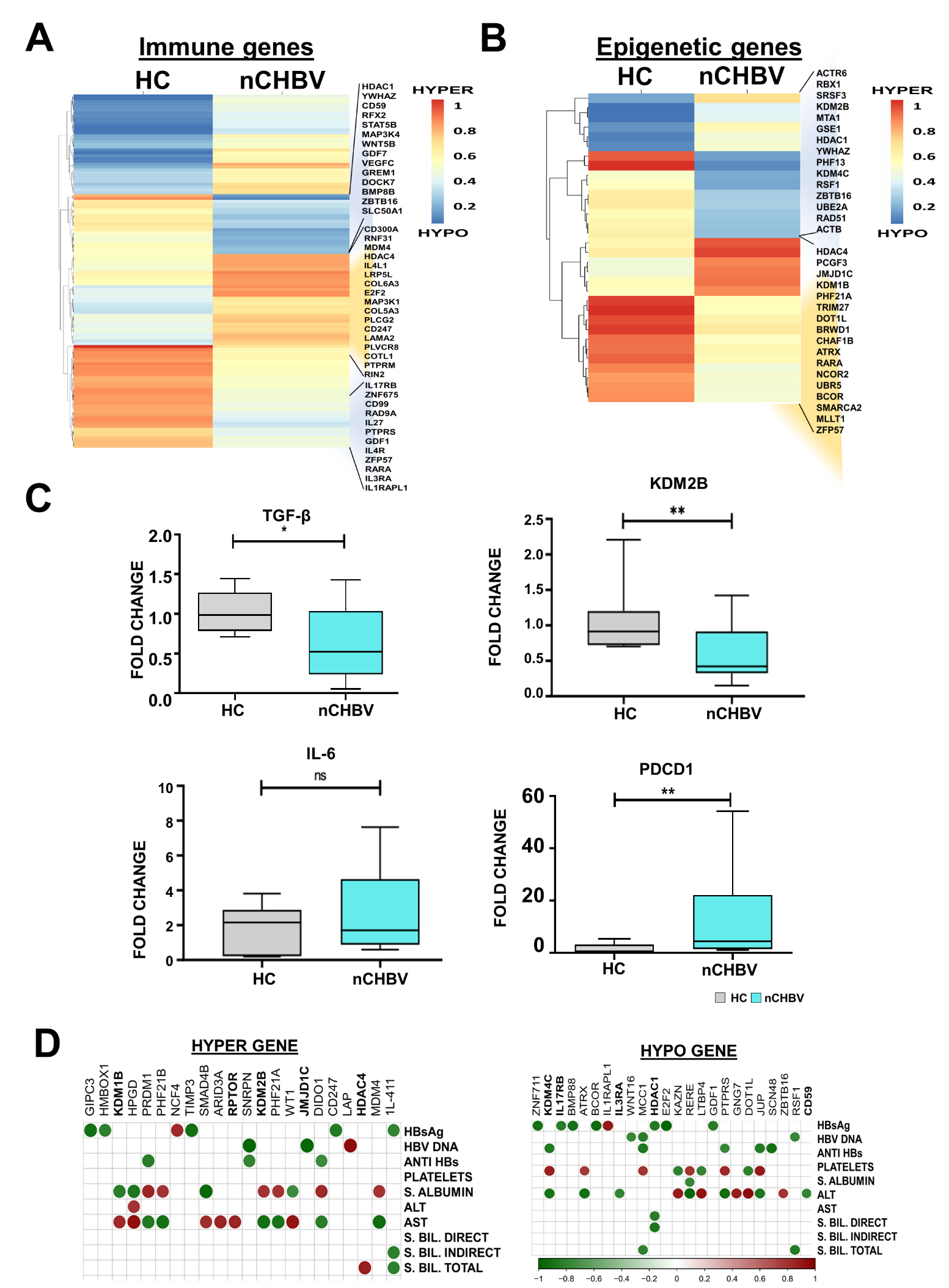

**Fig. S2.** **Altered methylation in nCHBV patients.** (A-B) Heatmap of differentially methylated immune and epigenetic genes depicting hypo and hypermethylation in nCHBV compared to HC. (C) Gene expression of KDM2B, TGF-β, IL6 and PDCD1 in PBMCs of nCHBV compared to HC (normalized with GAPDH). (D) Correlation analysis of significant hyper and hypomethylated genes with clinical parameters.

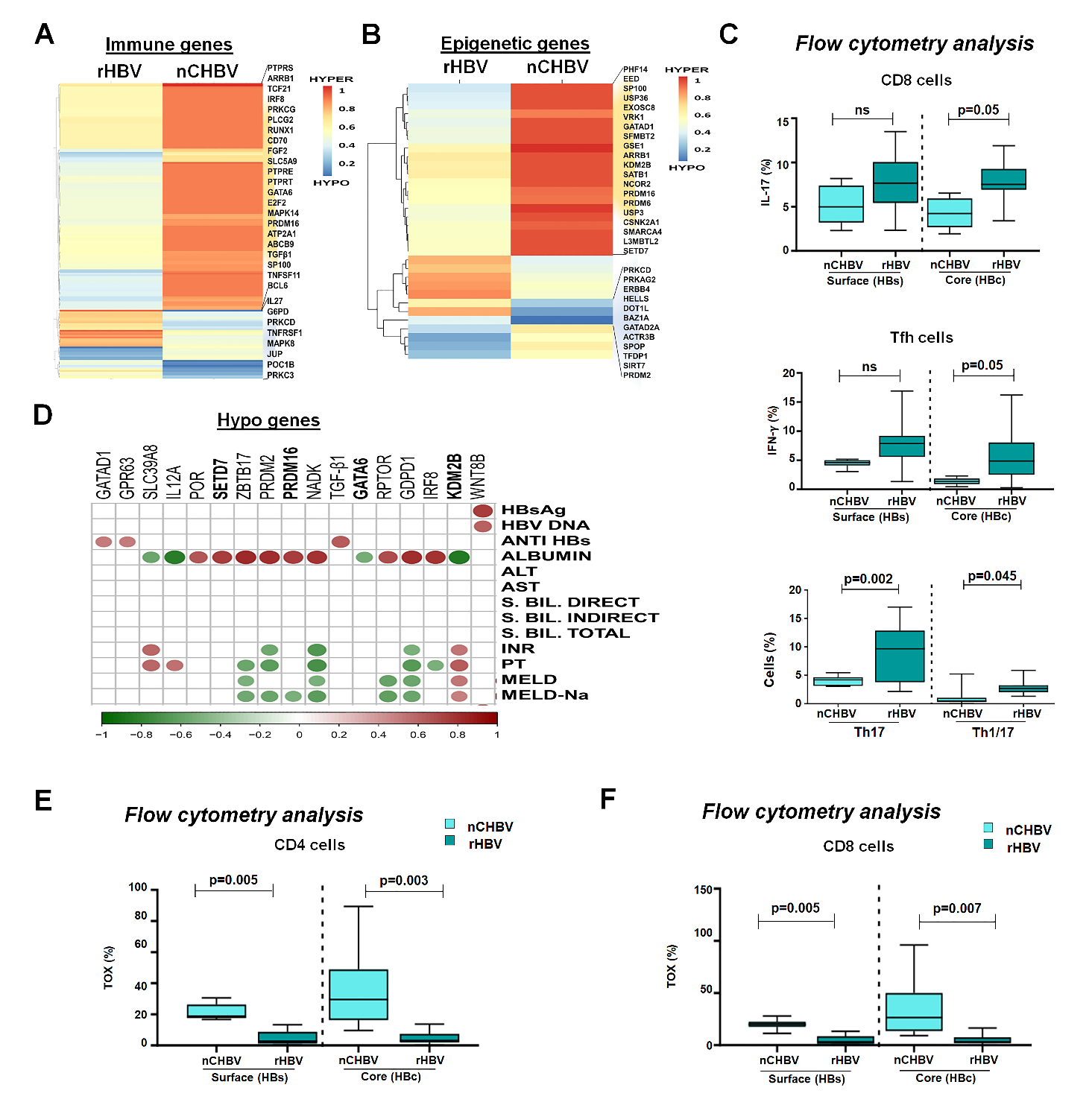

**Fig. S3.** **Reactivation suggests well defined epigenetic alteration leads to immune activation.** (A) Distribution of differential methylated Immune gene in rHBV and nCHBV. (B) Heatmaps of differentially methylated epigenetic genes in rHBV and nCHBV. (C) Percentage of IL-17, IFN-γ secreting HBV specific CD4, CD8, Th17 and Th1/17 cells in nCHBV and rHBV (D) Correlation analysis of significant hypo methylated genes with clinical parameters. (E-F) Expression of exhaustion marker TOX on HBV specific CD4 and CD8 T cells.

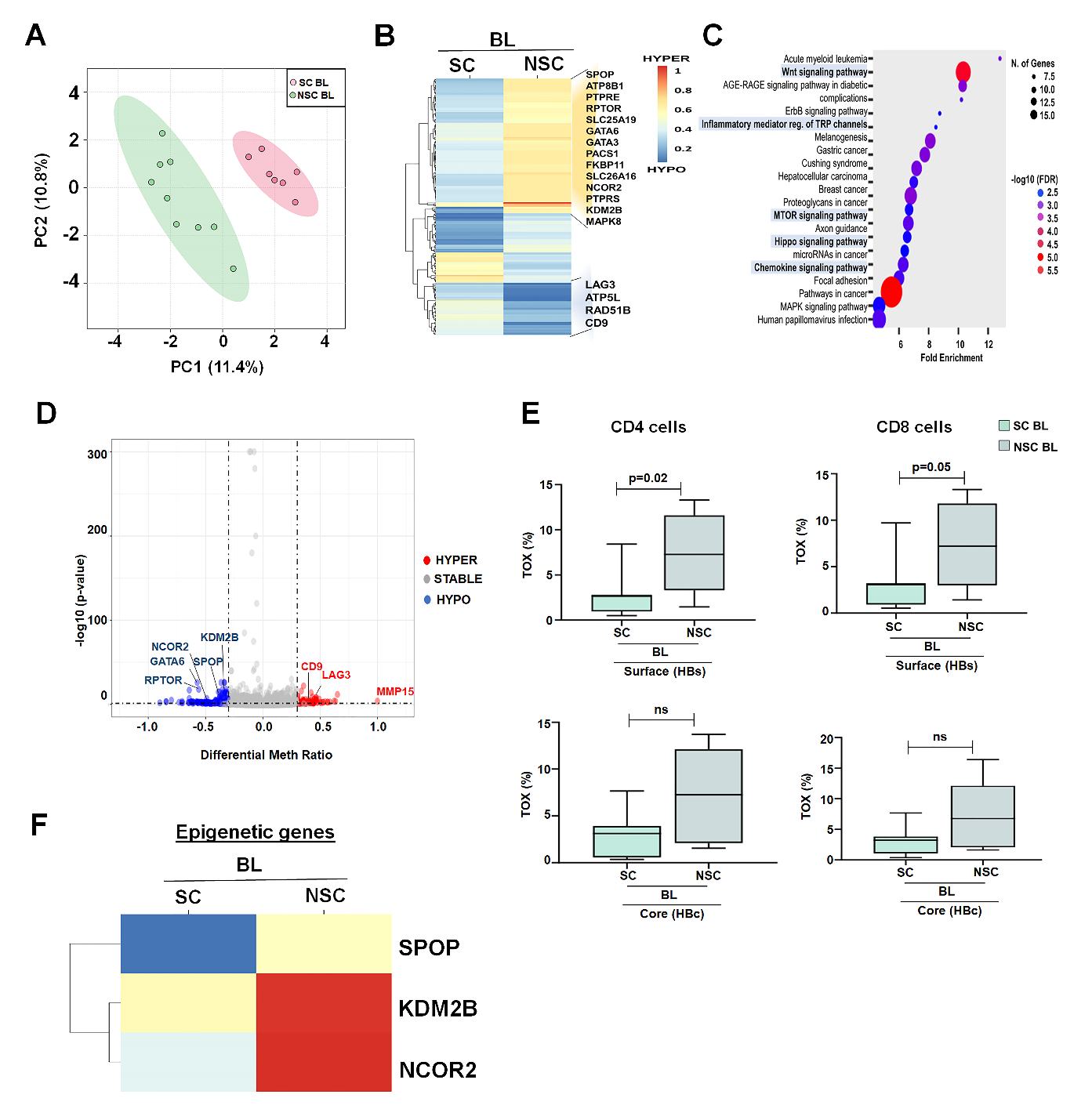

**Fig. S4.** **Immune dysfunction in non-seroconverters patients.** (A) Principal Component Analysis of DNA methylation in seroconverters (SC) and non-seroconverters (NSC) at baseline. (B) Heatmap showing unsupervised clustering of hyper and hypomethylated genes in SC and NSC at baseline. (C) GSEA depicts KEGG database pathway enrichment in SC patients compared to NSC. (D) Volcano plots show meth ratio (>0.33) vs -log10 p value (p<0.05) of selected genes in SC patients compared to NSC at baseline. (E) Expression distribution of exhaustion marker TOX on HBV specific CD4 and CD8 T cells. (F) Heatmaps of epigenetic remodellers represents hyper and hypomethylation in SC and NSC.

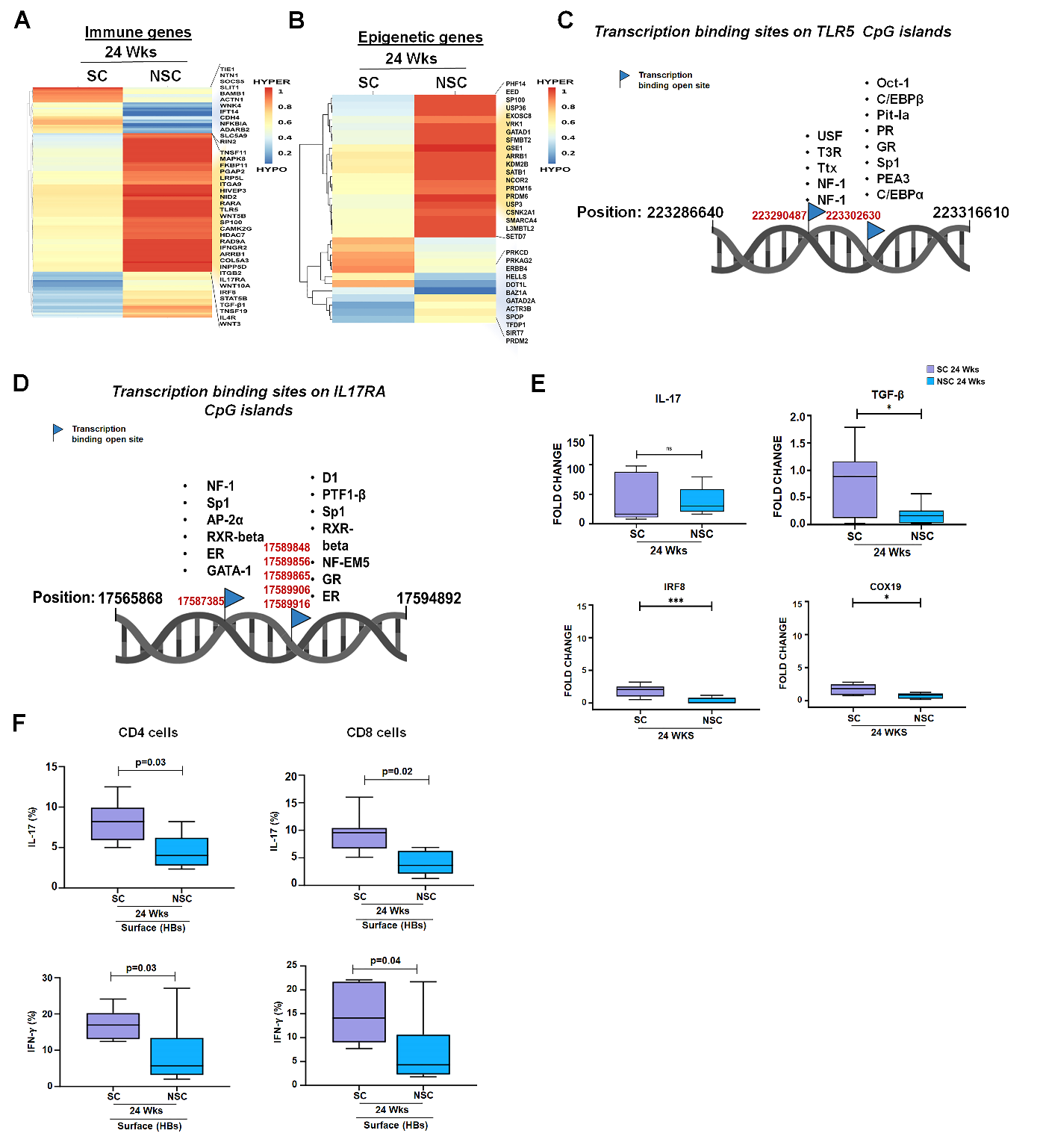

**Fig. S5. Chromatin accessibility of immune and metabolic genes at week 24.** (A) Heatmaps of immune genes depicts hyper and hypomethylated genes in seroconverters 24 wks and non-seroconverters 24 wks. (B) Heatmaps of epigenetic genes depicts hyper and hypomethylated genes in seroconverters 24 wks and non-seroconverters 24 wks. (C) TLR5 gene CpG contain transcription factor binding sites. (D) IL17RA gene CpG sites contain transcription factor binding sites. (E) Gene expression of IL-17, TGF-β, IRF8 and COX19 in SC and NSC patients at 24 wks. (F) IL-17 and IFN-γ secreting HBV specific CD4 and CD8 T cells.

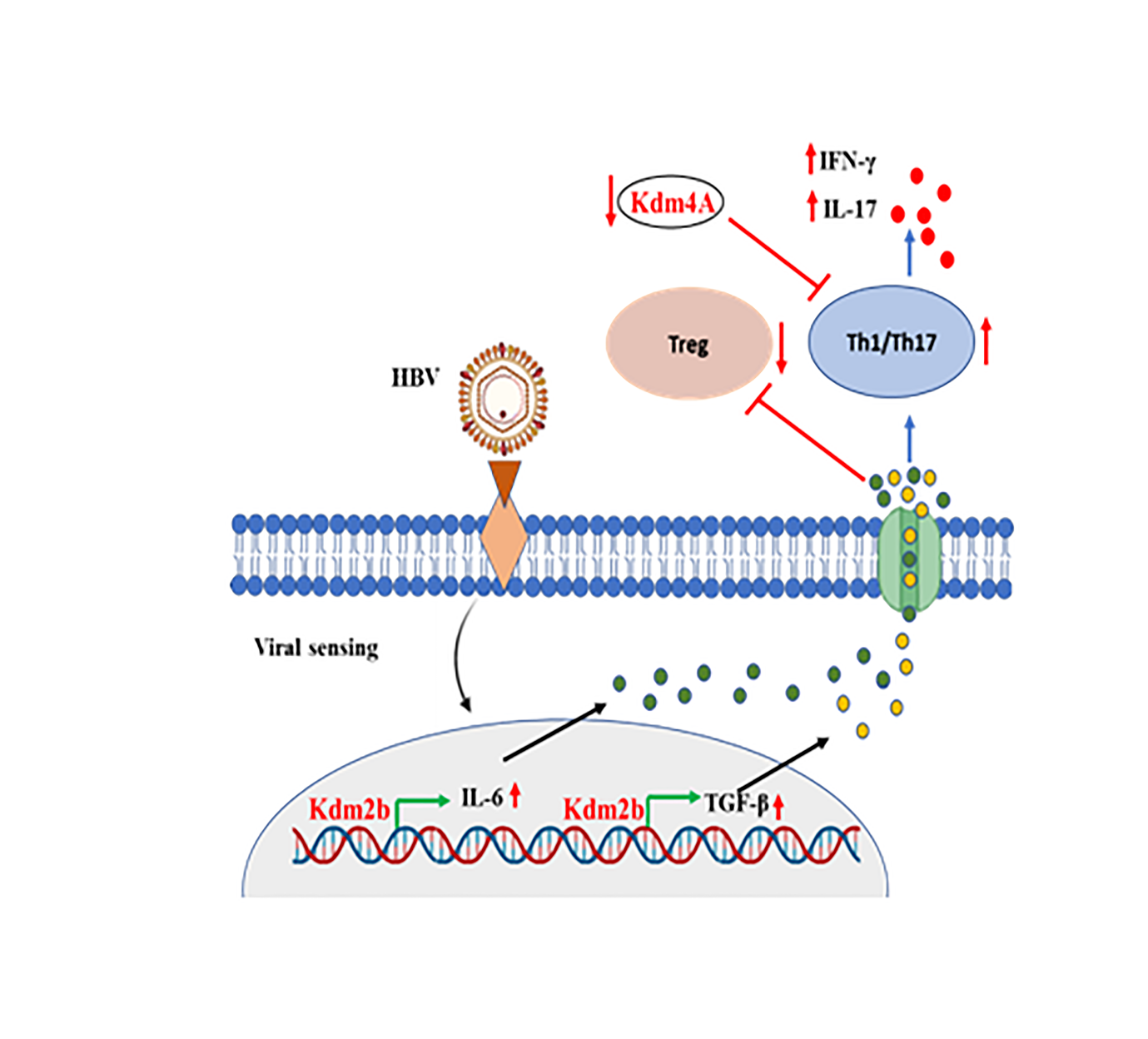

**Fig. S6**

**Supplementary Tables (see excel file)**

**Table S1.** Clinical and demographic characteristics of healthy subjects and HBV patients

| Variables | Healthy Control (n=7) | nCHBV (n=7) | HBV Reactivation (n=16) | *p values* | | |
| --- | --- | --- | --- | --- | --- | --- |
|  |  |  |  | HC  vs. nCHBV | HC  vs rHBV | nCHBV vs rHBV |
| Demographics | | | | | | |
| Age (years) | 28 (23-40) | 32 (22-62) | 35 (18-66) | 0.288 | 0.127 | 0.814 |
| Male:Female | 6/0 | 6/1 | 12/4 |  |  |  |
| Haematology | | | | | | |
| Haemoglobin (g/dL) | 15  (13-16) | 15 (14-16) | 14 (10-16) | 0.970 | 0.179 | 0.212 |
| Platelets (10^9/L) | 312 (278-376) | 207 (145-226) | 207 (81-377) | **0.002** | **0.001** | 0.499 |
| Leukocytes (10^9/L) | 7  (6-10) | 6 (5-6) | 7 (4-12) | **0.008** | 0.705 | **0.003** |
| Liver Function Test | | | | | | |
| S. Albumin (g/dL) | 4 (4-5) | 4 (4-5) | 4 (1-5) | 0.425 | **0.000** | **0.012** |
| ALT /SGPT (IU/L) | 30  (24-33) | 55 (25-116) | 357 (109-1862) | 0.055 | **0.000** | **0.001** |
| AST/SGOT (IU/L) | 32  (22-35) | 34 (22-141) | 315 (120-873) | 0.133 | **0.000** | **0.000** |
| S. Bilirubin Direct (mg/dl) | 0.1 (0.1-0.2) | 0.2 (0.1-0.2) | 6.3 (0.3-19.1) | **0.050** | **0.000** | **0.000** |
| S. Bilirubin Indirect (mg/dL) | 0.5 (0.2-0.8) | 0.9 (0.7-1.1) | 7.3 (0.5-22.2) | **0.005** | **0.000** | **0.000** |
| S. Bilirubin Total (mg/dl) | 0.7 (0.3-0.9) | 1.1 (0.9-1.6) | 12.2 (0.8-29.8) | **0.006** | **0.000** | **0.000** |
| S. Total Protein (g/dl) | 7.6 (7.3-7.9) | 7.8 (7.5-8) | 7.4 (4.4-8.7) | 0.071 | 0.099 | **0.003** |
| Kidney Function Test | | | | | | |
| S. Creatinine (mg/dL) | 0.79 (0.75-0.85) | 0.77 (0.76-0.77) | 0.7 (0.14-1.73) | 0.094 | 0.679 | 0.836 |
| S. Potassium (mEq/L) | 4 (4-5) | 4 (4-4) | 5 (3-5) | 0.705 | 0.259 | 0.213 |
| S. Sodium (mEq/L) | 138 (136-140) | 139 (137-140) | 136 (114-140) | 0.127 | 0.129 | **0.001** |
| Coagulopathy Test | | | | | | |
| INR | 0.6 (0.5-0.9) | 0.9 (0.9-1.4) | 1.2 (1-2) | 0.117 | **0.000** | 0.234 |
| PT (Seconds) | 12 (11-13) | 14 (14-14) | 15 (12-24) | **0.021** | **0.003** | 0.195 |
| Virological Parameters | | | | | | |
| HBsAg (IU/ml) | - | 21516 (2948-46013) | 6722 (250-125000) | - | **-** | 0.684 |
| HBeAg (n,%) | - | 2(28%) | 3 (19%) | **-** | - | - |
| HBV DNA (IU/ml) | - | 11000 (28-943000) | 31100 (2650-118000000) | - | - | 0.072 |
| Anti HBe (n, %) | - | 5(71%) | 13(81%) | - | - | - |
| Anti HBs (IU/ml) | 544 (256-987) | ND | ND | **-** | **-** | **-** |
| Anti HB core Total | - | 10 (9-10) | 10 (6-16) | **-** | **-** | 0.515 |
| Acute Complications | | | | | | |
| Ascites (%) | - | - | 3 | - | - | - |
| Hepatic Encephalopathy Grade 0 (%) | - | - | 14 | - | - | - |
| Hepatic Encephalopathy Grade I (%) | - | - | 0 | - | - | - |
| Hepatic Encephalopathy Grade II (%) | - | - | 2 | - | - | - |
| Hepatic Encephalopathy Grade III (%) | - | - | 0 | - | - | - |
| Liver Scores | | | | | | |
| MELD | - | - | 19 (6-31) | - | **-** | **-** |
| MELD-Na | - | - | 20 (9-34) | - | **-** | **-** |
| ACLF-CLIF | - | - | - | - | - | - |
| AARC-ACLF | - | - | - | - | - | - |
| SOFA | - | - | - | - | - | - |

**Table S2. Correlation of Methylated molecules with clinical parameters**

|  | **nCHBV** | |
| --- | --- | --- |
| ***Clinical Parameters*** | ***Positive*** | ***Negative*** |
| ALT |  | HDAC1 |
| AST | KDM1B | KDM2B  KDM4C  CD59  IL3RA |
| HBsAg | IL1RAPL1 | HDAC1  BCOR  IL17RB  ZNF711 |
| HBV DNA |  | JMJD1C |
| Albumin | KDM4C KDM2B | KDM1B  JMJD1C |
| Indirect Bilirubin |  | HDAC1 |

**Table S3.** Clinical parameters and methylated genes correlation in rHBV.

|  | **rHBV** | |
| --- | --- | --- |
|  | ***Significant Correlation*** | |
| ***Clinical Parameters*** | ***Positive*** | ***Negative*** |
| PT | KDM2B |  |
| INR | KDM2B |  |
| MELD | KDM2B |  |
| MELD(Na) | KDM2B |  |
| Albumin |  | GATA6  KDM2B |
| Platelets | PRDM16  SIRT7 | GATA6 |
| ALT/AST |  | IL27 |

**Table S4. Clinical and demographic characteristics of SC and NSC patients.**

| Variables | HBV REACTIVATION (n=16) | | |
| --- | --- | --- | --- |
|  | **SEROCONVERTERS (n=7)** | **NON-SEROCONVERTERS (n=9)** | **P values** |
| Demographics | | | |
| Age (years) | 35 (20-41) | 34 (18-66) | 0.251 |
| Male:Female | 5/2 | 7/2 |  |
| Haematology | | | |
| Haemoglobin (g/dl) | 14 (12-15) | 14 (10-16) | 0.810 |
| Platelets (10^9/L) | 250 (202-377) | 175 (81-334) | 0.072 |
| Leukocytes (10^9/L) | 8 (6-12) | 7 (4-11) | 0.366 |
| Liver Function Test | | | |
| S. Albumin (g/dl) | 4 (3-4) | 4 (1-5) | 0.911 |
| ALT /SGPT (IU/L) | 285 (238-1862) | 456 (109-1067) | 0.650 |
| AST/SGOT (IU/L) | 356 (189-859) | 266 (120-873) | 0.558 |
| S. Bilirubin Direct (mg/dl) | 10.1 (1.4-19.1) | 3.3 (0.3-10.5) | 0.092 |
| S. Bilirubin Indirect (mg/dl) | 7.3 (1.4-10.9) | 6.9 (0.5-22.2) | 0.769 |
| S. Bilirubin Total (mg/dl) | 18 (2.8-27.1) | 11 (0.8-29.8) | 0.346 |
| S. Total Protein (g/dl) | 7.3 (5.9-8) | 7.4 (4.4-8.7) | 0.892 |
| Kidney Function Test | | | |
| S. Creatinine (mg/dl) | 0.58 (0.14-0.92) | 0.88 (0.25-1.73) | 0.142 |
| S. Potassium (mEq/L) | 5 (3-5) | 5 (4-5) | 0.519 |
| S. Sodium (mEq/L) | 136 (131-138) | 137 (114-140) | 0.690 |
| Coagulopathy Test | | | |
| INR | 1.2 (1-2) | 1.1 (1-1.9) | 0.527 |
| PT (Seconds) | 15 (12-23) | 15 (12-24) | 0.814 |
| Virological Parameters | | | |
| HBsAg (IU/ml) | 9627 (1266-28967) | 5263 (250-125000) | 0.640 |
| HBeAg (n,%) | 2 (29%) | 1 (11%) | 0.225 |
| HBV DNA (IU/ml) | 31100 (3250-7690000) | 1515500 (2650-118000000) | 0.089 |
| Anti Hbe (n,%) | 5 (71%) | 8 (89%) | 0.628 |
| Anti HBs (IU/ml) | 0 (0-0) | 0 (0-0) | 0.437 |
| Anti HB core Total | 10 (8-23) | 10 (0-26) | 0.947 |
| Acute Complications | | | |
| Ascites (%) | 1 | 2 | - |
| Hepatic Encephalopathy Grade 0 (%) | 7 | 7 | - |
| Hepatic Encephalopathy Grade I (%) | 0 | 0 | - |
| Hepatic Encephalopathy Grade II (%) | 0 | 2 | - |
| Hepatic Encephalopathy Grade III (%) | 0 | 0 | - |
| Liver Scores | | | |
| MELD | 20 (12-29) | 15 (6-31) | 0.280 |
| MELD-Na | 22 (15-32) | 19 (9-34) | 0.394 |
| ACLF-CLIF | - | - | - |
| AARC-ACLF | - | - | - |
| SOFA | - | - | - |
